# User-friendly high-content imaging analysis on a single desktop: R package H5CellProfiler

**DOI:** 10.1101/2022.10.06.511212

**Authors:** Steven Wink, Gerhard A. Burger, Sylvia E. Le Dévédec, Joost B. Beltman, Bob van de Water

**Author notes:** **Corresponding Author** Bob van de Water, GORLAEUS BUILDING, EINSTEINWEG 55, 2333 CC LEIDEN, THE NETHERLANDS, *E-mail address:.

## Abstract

Technological development has led to ever-increasing amounts of data in high-content screening. For utilizing such data in an efficient, thorough, and user-friendly manner we developed the R package H5CellProfiler. H5CellProfiler is based on R packages data.table, ggvis, ggplot2 and shiny. H5CellProfiler launches a browser that allows scientists to analyze large single-cell datasets and make statistical summaries and graphs on their local desktop in a fast and memory-efficient manner. In addition, single-cell track labels are calculated and broken tracks are re-connected based on user-defined thresholds resulting in unique sets of annotated tracks.

## Introduction

High-throughput high-content automated imaging (HT-HCI) includes image acquisition, image storage, image annotation, image analysis, data analysis, and storage of the analysis results. Image acquisition is performed by high-end automated microscopes that are equipped with hardware and software for stage movement or objective movement and often with an incubation chamber that has temperature and CO2 regulation allowing live cell imaging. These conditions allow the acquisition of large imaging datasets from multi-well plates capturing the dynamics of cell biological events.

Image annotation can be divided into two categories: first, technical annotation which is automatically generated by the acquisition software and consists of technical acquisition parameters. Second, biological annotation consisting of parameters that the scientist needs to add himself. The biological annotation of images is closely related to the data storage of images in a file server, which is managed by a database management system (DBMS). The DBMS allows annotation, organization, querying, and often visualization of the acquired images. Well-known image DBM systems are OMERO from the Open Microscopy Environment (OME) (Allan et al., 2012) and its commercialized version Columbus.

Image analysis can be performed with the aid of commercial or open-source software which differ in the offered functionality and flexibility. Several commercial software systems exist that provide easy-to-use image analysis solutions, sometimes directly combined with image acquisition. Well-known software systems include NIS-Elements (NIKON), Metamorph (Molecular Devices), AxioVision (Zeiss), and Volocity (Perkin-Elmer). Recently, commercial systems have been developed that even combine multiple HT-HCI stages. For example, Columbus from PerkinElmer can store, annotate, and analyze images and analyze the resultant data with several statistical and graphical tools due to integration with TIBCO Spotfire.

A disadvantage of using commercial software for image analysis is that it is difficult or simply not possible for researchers to modify or extend it because the source code and even the applied image analysis algorithms are hidden to secure the commercial product. Commercial systems are therefore best suited for standard image analysis pipelines and are mostly used by industry or by labs with relatively basic and fixed image analysis needs.

Labs that frequently change equipment or have specialized image analysis requirements often prefer open-source solutions. The most well-known solution is ImageJ (Schindelin et al., 2012), which is an important platform for scientific image analysis development. However, it can be challenging to build workflows for automated image analysis because it requires some knowledge of the available ImageJ plugin libraries, a decent understanding of image analysis algorithms, and Java scripting skills. For this reason, some researchers prefer user-friendly, GUI-based image analysis software tools. A well-known example is CellProfiler (Kamentsky et al., 2011), which was developed initially by Carpenter and colleagues. In CellProfiler, parameter optimization can be performed with the help of the GUI and accompanying graphical displays of the segmentation results. Subsequently, image analysis with appropriate parameters can be performed in an automated fashion. In parallel with recent technological advancements in microscopy, the interest in the analysis of single-cell behavior has strongly increased. Indeed, research on the heterogeneity of cell populations (Forslund et al., 2012; Pauwels et al., 2012), the morphology of subcellular structures (Di et al., 2014), transcription factor oscillatory dynamics (Herpers et al., 2016) or cell migration (Roosmalen et al., 2015) all require single-cell measurements. As a result, the amount of data has strongly increased with consequences for the handling of image analysis output. A typical overnight live single-cell imaging session of an in vitro 2D 384-well plate leads to around 20 GB of raw imaging data (approximately 20,000 images) and depending on the features of interest up to 5 GB of analysis data. Memory limitations require data dumps during the analysis by temporary storage in relational databases or hierarchical file-based formats. Hence, labs need to set up a dedicated database server which requires maintenance and technical expertise, which can be challenging for biology-oriented labs.

CellProfiler 2 and 3 include a feature for storing measurements during the analysis using the file format HDF5 (Fraser, 2015), a portable data format without size limitations due to the hierarchical structure. An HDF5 file can be navigated through the use of file paths – strings separated by the forward slash (/) to define the hierarchical structure of the data, analogous to the Unix file system. In addition, the HDF5 format has been implemented specifically for cell-based assays in high-content microscopy in the form of CellH5 which can be used as a format for data exchange. CellH5 and several CellH5 interfaces have been developed by Huber and colleagues (Sommer et al., 2013), allowing for visualization of the stored images and reading and writing of the quantitative data. However, for CellProfiler HDF5 output user-friendly solutions do not yet exist. Therefore, we have developed the R package H5CellProfiler. H5CellProfiler requires a metadata file with biological annotations and an HDF5 file with the quantitative measurements from CellProfiler. The package loads the data into memory and organizes user-requested features at the level of single cells. Subsequently, a browser is launched and functions as a GUI for the user to manipulate, summarize and display the single-cell data. This enables biologists to perform their image analysis with CellProfiler followed by the semi-automated production of single-cell data summaries and visualizations that would not be possible using spreadsheets. Thus, H5CellProfiler allows to perform all data operations on a single desktop without the need to set up database and client software and as such makes the entire image analysis workflow user-friendly (Fig. 1).

**Figure 1:**
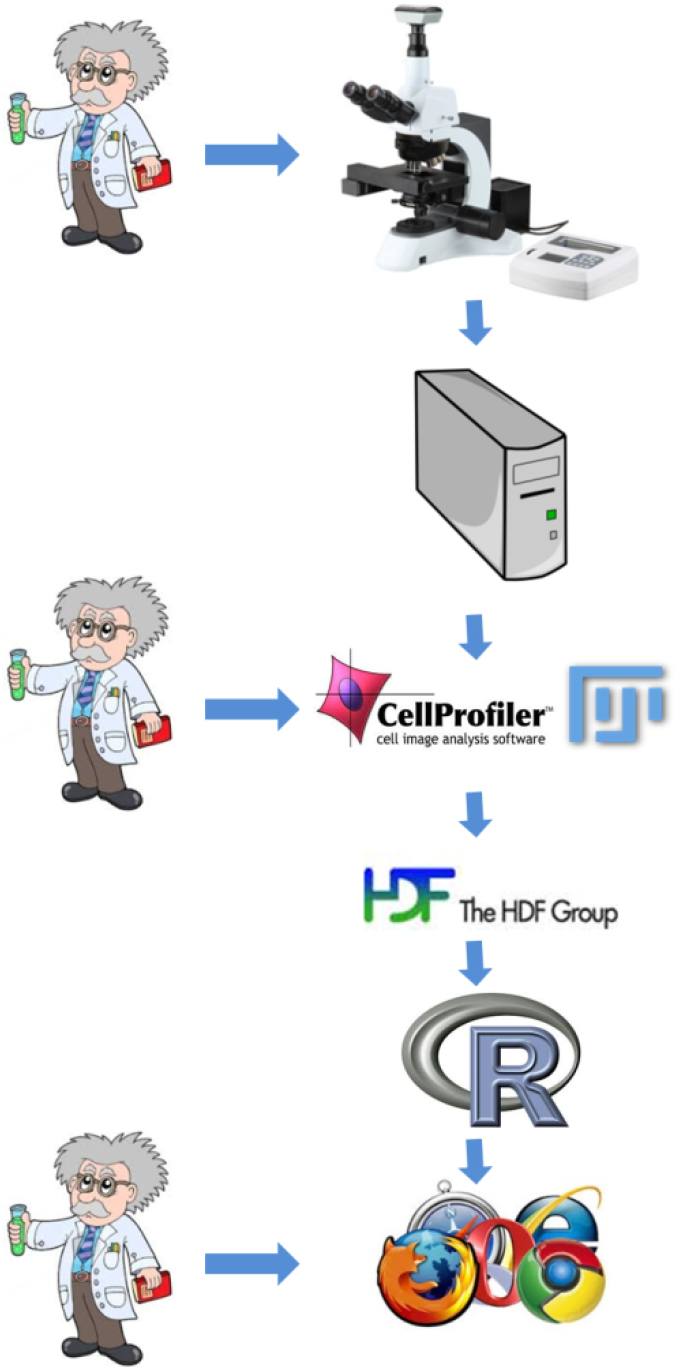
User-friendly High Content image analysis workflow. The scientist interacts with the imaging device, sets up the image analysis with GUI-based CellProfiler run on the local desktop, and finally performs the data analysis with a browser-based GUI which communicates with the R server (usually on the local desktop).

## Results

### Description of the H5CellProfiler package

#### H5CellProfiler architecture

H5CellProfiler reads the HDF5 data and uses the input arguments (which can be typed in the R terminal or provided as a file) and annotation files provided by the user to reformat and summarize the data in a flexible, intelligent, and computationally efficient manner (Fig. 2). It is flexible because the user guides the reformatting with help of the GUI and has several options: extracting relevant data into a large text file containing a single row per parent object, making a summary for each treatment, writing a text file containing rows per tracked object and columns corresponding to the time points for each time-lapse or plotting graphical displays of the data The size of the data hardly affects the performance as long as it does not exceed the available memory capacity. This is because all data manipulations are very fast due to parallel computing and ordered indexing. As a result, up to 100GB of data is easily handled by H5CellProfiler.

**Figure 2:**
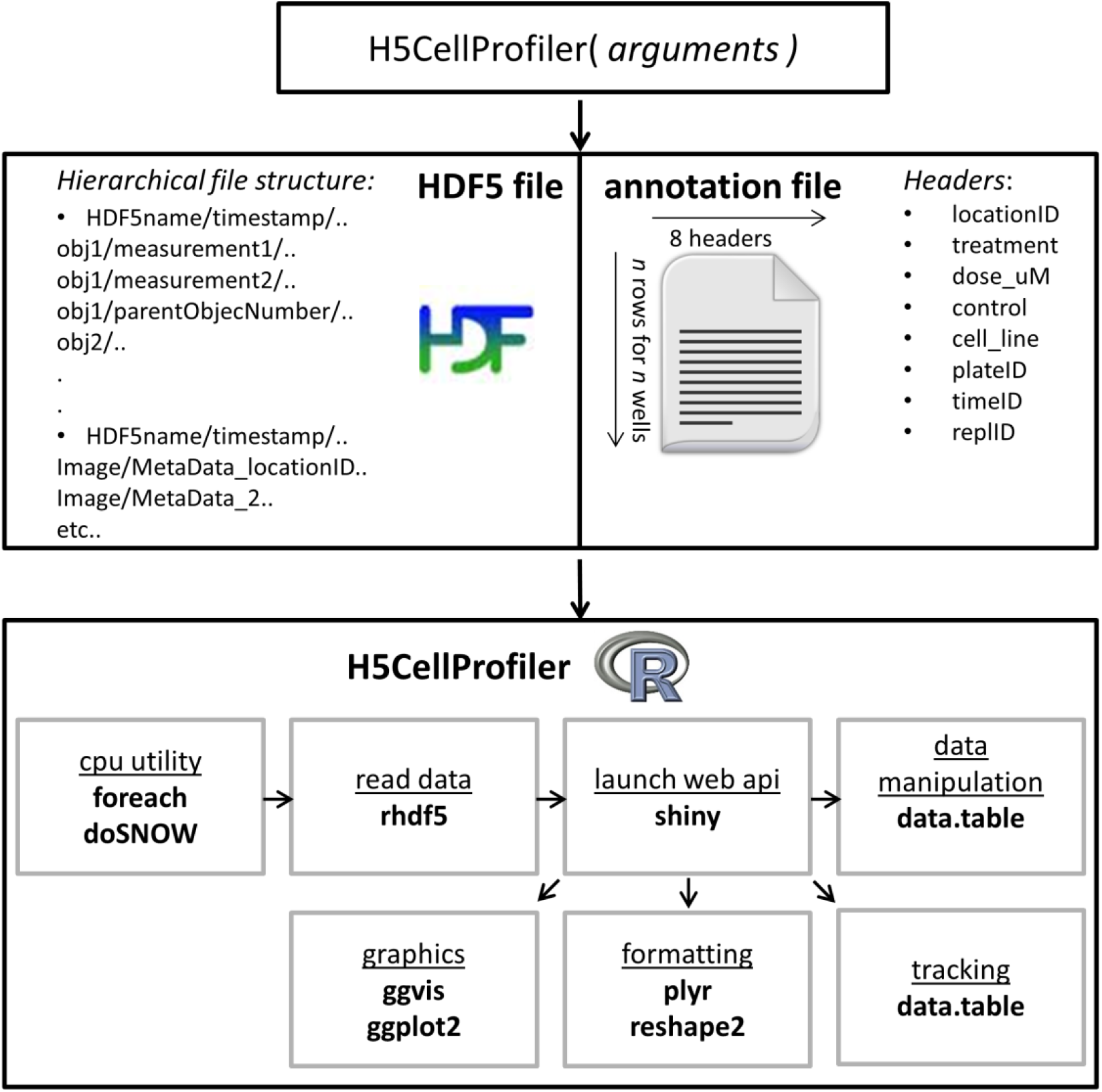
Architecture of H5CellProfiler. The main function is run by typing the H5CellProfiler (*arguments*) function in the terminal or by providing the arguments in a file. H5CellProfiler then uses the HDF5 file containing the image analysis data and the annotation file provided by the user containing the biological annotation. The architecture of the H5CellProfiler package is depicted in the lower panel: the order of the separate internal processes (underlined) using various R packages (**bold**) are depicted by the arrows.

#### Parallel computing

H5CellProfiler offers parallel processing to speed up the analysis of large datasets. This is achieved with the parallel computing package ‘foreach’ and parallel back-end registration for windows operating systems ‘doSNOW’. The number of parallel sessions that can be initiated for reading and data analysis is set by the user with a maximum equal to the number of separate HDF5 files. Parallel computing especially speeds up the summary calculations, the data reshaping for plotting, and the re-labeling of tracking data (the number of cores for re-labeling the tracking data is independent of the number of HDF5 files and is set by the user-set argument). Chunks of the data in memory are divided over the available cores without duplication of the data for each core, thus ensuring memory-efficient parallel computing.

#### Reading HDF5 and formatting data

We employed the R package ‘rhdf5’ (Sommer et al., 2013) for reading and writing HDF5 files. An advantage of using the HDF5 format to store and read data is its annotated hierarchical hyper-rectangular structure. Together with the annotation, this enables the selection of data subsets without the need to query through the entire dataset. As a consequence, very large datasets do not slow down reading significantly.

### Application of R package H5CellProfiler

Basic knowledge of CellProfiler is required to be able to provide the correct arguments for the H5CellProfiler function and are therefore briefly described here. A detailed manual of CellProfiler can be found on the website https://cellprofiler.org/.

#### Regular expressions that enable linkage of biological annotation with image data

In CellProfiler, variables are captured using a simple collection of regular expressions. For example, for an image named `xy0013c1t01`.tif and located in a folder named `../C03` (according to the well number in the plate), two sets of regular expressions capture this image annotation. In this case, the first regular expression would be `(?P<imageNumber>xy[0-9]{4})c[1-4]{1}(?P<tp>t[0-9]{2}).tif` to capture the `image number` 0013 in the CellProfiler variable imageNumber and the time point 01 in the variable tp. Note that the characters inside the square brackets are the characters to be searched for in the image file names, the numbers inside the curly brackets denote the number of such characters, and the construction with `(?P<variable_name>…)` determines in which CellProfiler variable the contents will be stored. Similarly, the folder name ../C03 can be captured by the regular expression `/(?P<Well>.*)$` or by `/(?P<Well>[A-Z]{1}[0-9]{2})$` in the CellProfiler variable `Well`. These expressions search well information between the last forward slash and the end of the string ($), for example, `C03` in the string `../C03`. The first expression is flexible, the second is more constrained in the formats it accepts. For more information, see https://regexr.com/. The key feature in these examples is that images are automatically annotated by CellProfiler and that this information is stored together with the image analysis results in the HDF5 file. The annotations obtained by the regular expressions are stored in the HDF5 output and the HDF5 file path is defined as ‘Metadata *variable name*’ Together with a user-provided annotation text file, this enables H5CellProfiler to link biological annotation to the image measurements (Fig. 2). A tab-delimited text file containing the biological annotation is created by the scientist which consists of several columns/headers and a row for each well of all multi-well plates that are simultaneously analyzed with H5CellProfiler. The image analysis results of separate plates can be stored in a single HDF5 or multiple HDF5 files, depending on how the images were analyzed. In addition to the well (column ‘locationID’) and plate id (column ‘plateID’) several optional columns can be entered: the column ‘treatment’ can be used to enter annotation regarding, for example, compound or siRNA treatment of the well. Moreover, if the column ‘dose μM’ is used to annotate the applied concentrations, dose-response curves can be automatically generated at a later stage using the GUI. Furthermore, it is possible to use columns to annotate replicate numbers (‘replID’) and time information (‘timeID’) which can be useful for imaging of multiple plates that represent different time points. Finally, the column ‘control’ can be used to enter control information, which can be useful for plate normalization purposes in the GUI.

#### H5CellProfiler arguments

The H5CellProfiler package requires several of these meta-data HDF5 file paths as arguments to the H5CellProfiler function. At least `locationID` and `imageID` are required, and if multiple plates have been stored in a single HDF5 file also `plateID` must be provided as an HDF5 file path. In addition, it is possible to provide `timeID` and `replicateID` if required. The `plateID`, `timeID`, and `replID` arguments can be defined either as an HDF5 file path or as a constant for a single or for multiple HDF5 files (the latter is only possible if these variables remain the same within a single HDF5 file.

Conversion of the integer values for the time (either by regular expression or manually defined per HDF5 file) to real experimental time is performed based on the two variables `exposureDelay` and `timeBetweenFrames`. These represent the delay of the first image relative to the biological perturbation of interest and the time between two consecutive time points, respectively.

The CellProfiler nomenclature consists of objects and measurements on these objects. Objects can be cells, parts of cells (e.g., cytoplasm) but also the images themselves. Users will segment the objects of interest and define the name of these objects. For each object, several measurement categories can be chosen in CellProfiler, for example, intensity, texture, and morphological measurements can be passed on to H5CellProfiler. Up to ten of these object measurements can be passed on to H5CellProfiler in the `myFeaturePathsA` variable. The single entries of these arguments should have an identical format as the CellProfiler output namely ‘object/MeasurementCategory measurement’.

Objects are often related to other objects. For example, when there are multiple objects for each cell, a parent object must be defined to link the secondary objects to. In this manner, every object defined by segmentation or object manipulations such as subtraction or shrinking has an object number and an associated parent object number. These object relations are provided to H5CellProfiler as the variables `parentObject, childObject1, childObject2, …, childObject5` and `tertiaryObject`, thus defining how the single-cell data are organized. The `tertiaryObject` is a CellProfiler-defined child object defined by other-object manipulations such as subtraction (often the cytosol, defined as the cell subtracted by the nucleus). The image and parent object define the primary key of the tables in memory and secondary objects are associated with their corresponding parent object and thus on the same row in the table. Thus each row in the table corresponds to a unique parent object. If multiple secondary objects exist for single parent object (e.g., multiple organelles belonging to a single parent nucleus), a summary statistic must be calculated. This summary statistic is by default the mean but can be added as an argument in the form of an R function. Recommended statistics include the mean or a particular quantile, and are defined as a function, for example, `function(x) { mean(x, na.rm = TRUE) }` used as the `multiplePerParentFunction` variable.

Finally, the number of CPU cores the user wants to assign for all data processing steps can also be provided to H5CellProfiler as the numberCores variable. However this is limited to maximally the number of HDF5 files provided, however, for the tracking optimization algorithms this is only limited by the number of individual locations that were imaged (number of time-lapses)

#### Track re-labeling and optimization

CellProfiler also performs tracking in the context of time-lapse imaging, which means that objects are linked to each other in time. The H5CellProfiler package recognizes tracking parameters automatically using regular expressions, so the user does not need to provide additional arguments. However, the `fixTrackingFunction` utility from H5CellProfiler offers several options like re-labeling and combining ‘broken’ tracks based on user input. During tracking with CellProfiler, segmentation errors frequently lead to track interruption for one or several frames. These errors are quite common when cells tend to cluster together while migrating, resulting in an underestimation of the number of segmented objects. Alternatively, overestimation of segmented objects can also cause tracks to break because the tracking label can connect to the spurious cell which will disappear in the next few frames. Tracking errors are maintained in time, meaning that a small percentage of segmentation errors per time point results in an accumulation of errors in track labeling. CellProfiler labels a tracked cell based on the object it is estimated to originate from at the previous time step and can assign the same label to multiple objects. Thus, a cell division or a single frame segmentation error results in the same label being assigned to multiple tracks. The `fixTrackingFunction` function re-labels these to new tracks starting at the last time point of imaging, using the track-parent object labels assigned by CellProfiler. A track parent and track child are related through time, thus corresponding to the same tracked object. When multiple track-child objects point to an identical track-parent object (in the previous time point) the track label will be assigned to the closest parent using the x-y coordinates of the parent and child objects. After the re-labeling, the algorithm additionally attempts to reconnect broken tracks over 1-3 time frames if the parameter `reconnect_frames` is set by the user. In addition, the maximum pixel distance to reconnect partial tracks can be set with the parameters `max_pixel_reconnect1, max_pixel_reconnect2` and `max_pixel_reconnect3`. If multiple tracks are within the distance threshold the closest tracks are connected. Finally, the minimal track length can be set with the parameter `minTrackedFrames` which means that only tracks with at least that length after relabeling and reconnection are retained. The tracks can be plotted using the GUI and it is up to the user to validate the reconnect-argument settings by comparing the track plots with the time lapses.

### Graphical User Interface

After the selected data has been read into memory and is reorganized with the help of the parent-child object relations and the annotation file, a browser is launched (Fig. 3). This utility is based on the package ‘shiny’, which is an R wrapper for HTML and JavaScript. Shiny requires a `server` and `ui` definition. The ui defines the layout and functionality of the GUI and generates the tools required for user input and output. The server defines the R `server` that performs the operations. These definitions can be provided as separate `ui.r` and `server.r` files, but can also be combined in an `app.r` file.

**Figure 3:**
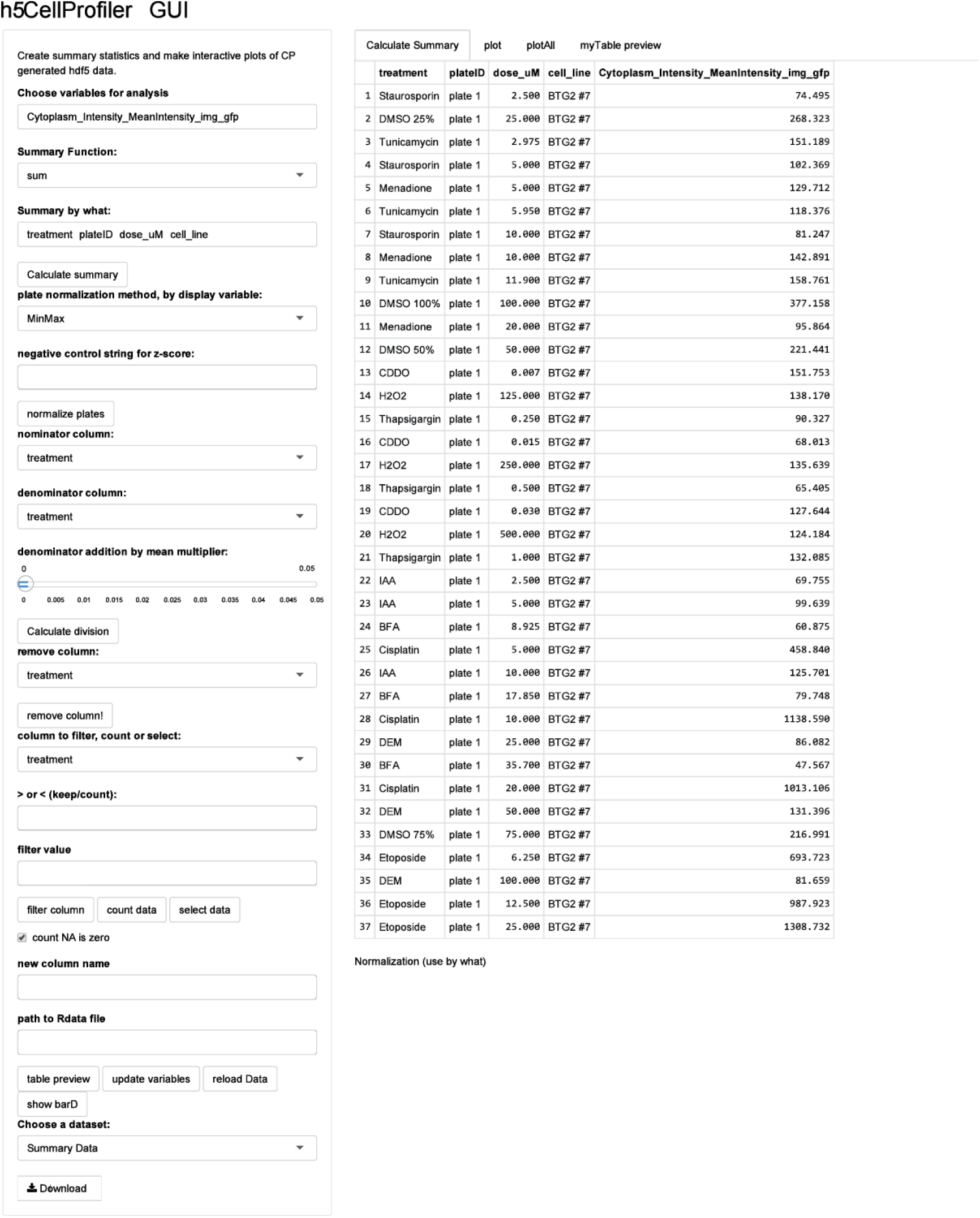
GUI for HCI data. Left panel shows an example of a GUI data manipulation and summarization utility page. Utilities include summarization statistics over the requested variables and factors. Further options include choice of plate normalization methods, column division, column deletion, filtering, selection, counting and downloading the modified single-cell data or summarized data. Right panel shows an example of a summarized table preview within the browser.

The web application provides a graphical user interface for frequently used data manipulation, summarization, and graphical operations. It sends commands to an R session so the user only indirectly employs R commands. This approach helps to ensure that the data manipulations are performed correctly.

The user can choose one or multiple variables for summarization, and one or multiple factors over which the summarization takes place. The offered summary functions are the mean, standard deviation, median, sum, minimum, maximum, and several quantiles. As soon as these options have been selected the table is calculated using the R package ‘data.table’ and takes fractions of a second for tables up to 1GB. The raw measurement variables can also be normalized by z-score, by the plate median, or by the min-max method (*x − x*_plate min_)*/*(*x*_plate_max_ *− x*_plate_min_). These normalizations are performed over the summarized results as defined by the `by.what` factors because single-cell data contain too many extreme outliers.

Other data manipulations provided by the browser include column division, deletions, filtering, counting, and selection. Finally, the modified single-cell table and summarized table can both be downloaded as tab-delimited text files. When the summarized table has been created the ‘plot’ tab (Fig. 3, right panel) provides the option to explore the data graphically in an interactive manner (Fig. 4). The user can select the treatment to be displayed and the variables to be plotted. Moreover, a third dimension in the data can be visualized with color mapping.

**Figure 4:**
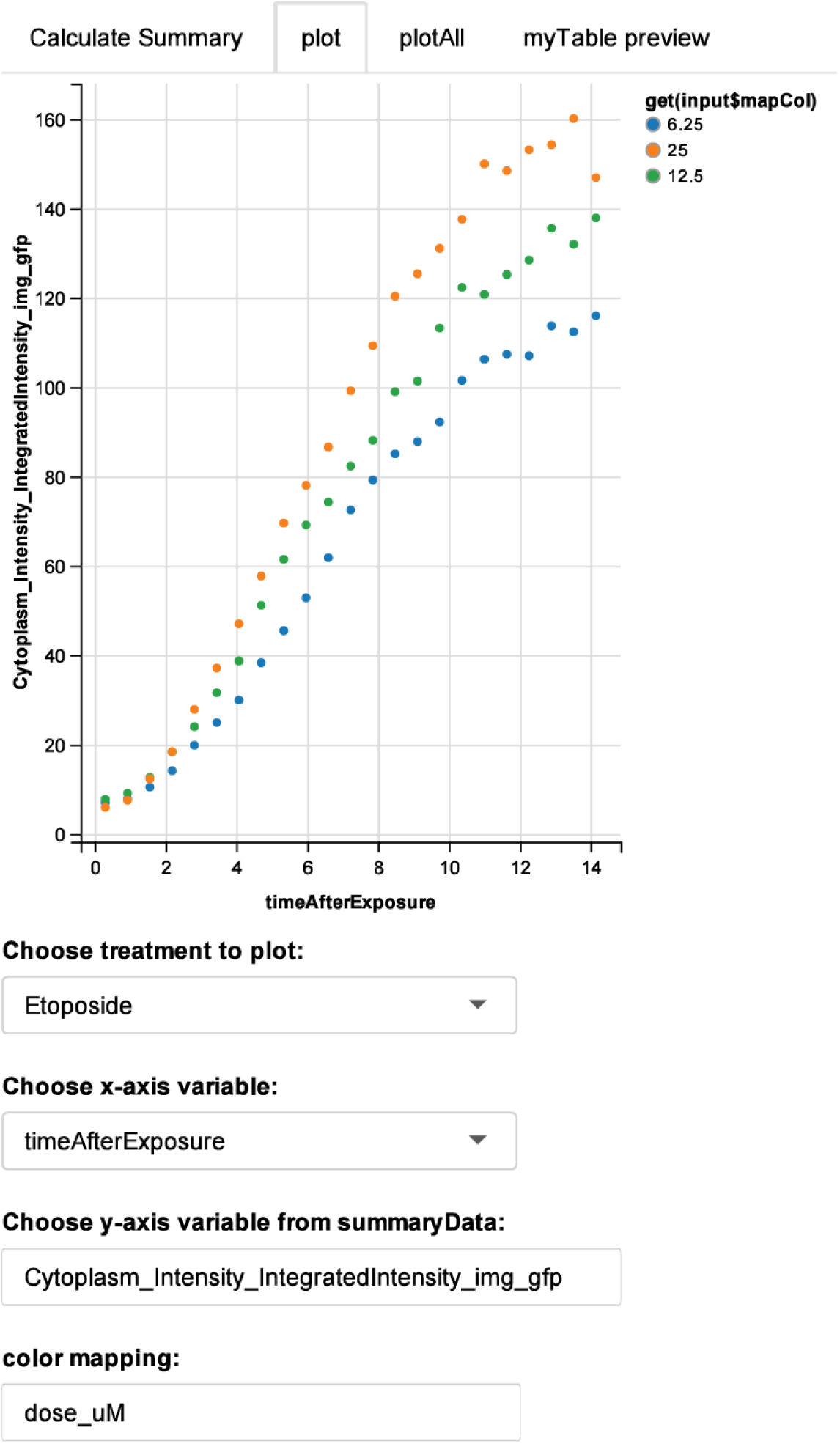
Interactive plotting with help of the package ‘ggvis’. A treatment, x-and y-axis variable and a color mapping can be selected from a drop down box, after which the plot will automatically update.

The R package ‘ggplot2’ offers faceting of variables which means that multiple graphs can be automatically wrapped in a single figure. This option to plot large numbers of graphs at once is offered by the ‘plotAll’ tab (Fig. 4, right panel). In Fig. 5, an example of graph faceting is shown, together with the input boxes. With this approach, it is possible to plot hundreds of graphs in a single figure if the PDF size is sufficiently large for the graph annotations to be displayed correctly (with a small font). Options for multi-faceted plotting include the plot type (line plot or bar plots), variable collapse (i.e., calculation of the mean), and the choice of x and y variables. Moreover, the user can map up to six variables or specific values by employing different color, fill, shape, and size aesthetics. Other options include adding lines between the dots and setting the PDF size and font size. The minimum and maximum values of the x- and y-axis can be set to ‘fixed’ (i.e. employing the minimum and maximum values of the entire dataset), to ‘free’ (i.e. specific ranges for each plot), or can be set manually. Finally, the dose values can be set to ‘dose-levels’ instead of their specific dose ranges which are usually different for each treatment (see example in Fig. 5). Dose level 1 is the lowest value in a dose range and the highest dose level corresponds to the highest concentration employed for each treatment separately.

**Figure 5:**
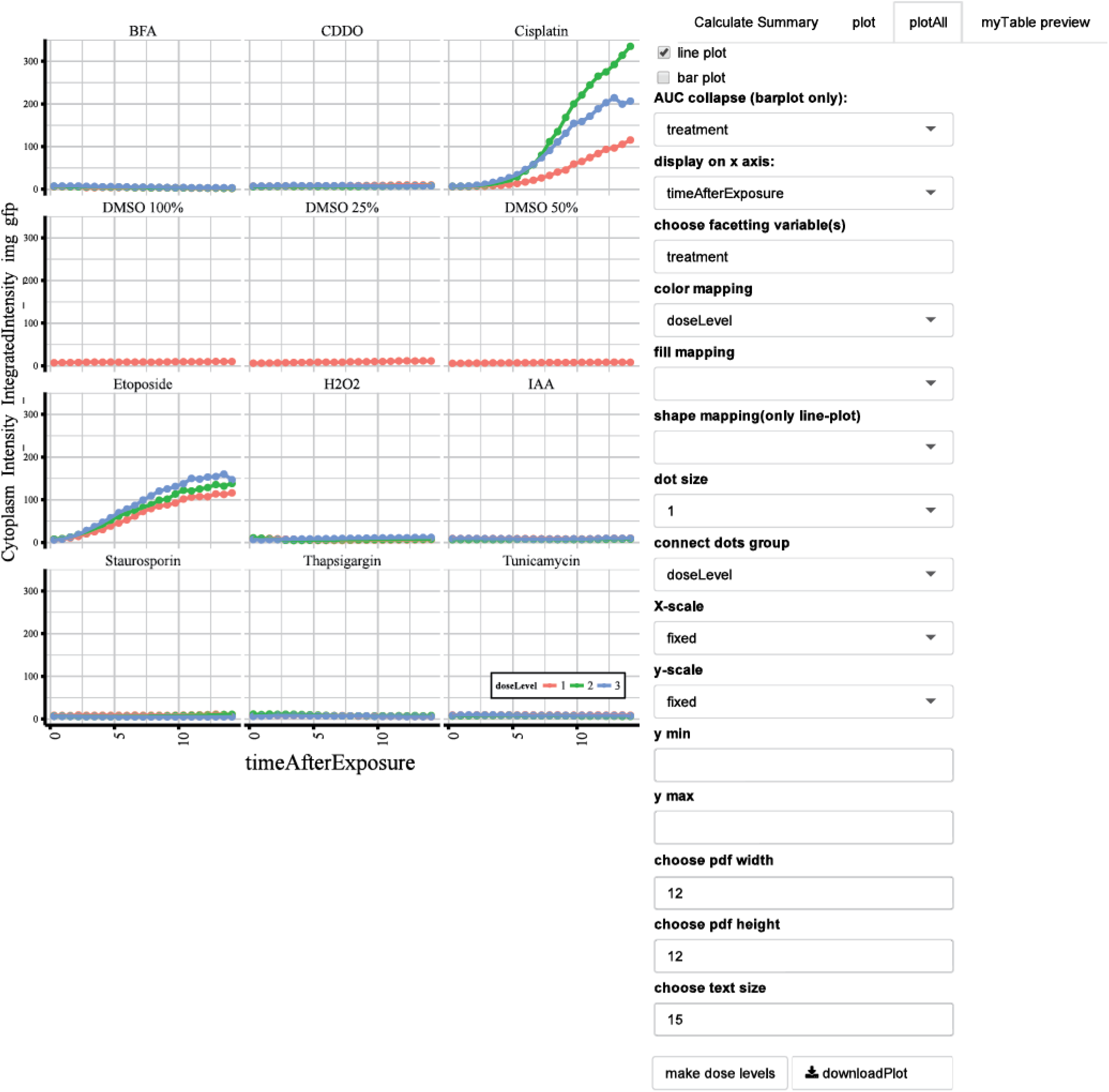
Multi-faceted plotting with help of package ggplot2. In the ‘plotAll’ tab the specifics of faceted plotting can be defined. The options are provided in the browser as displayed in the right panel.

The tab ‘myTable preview’ (Fig. 3, right panel) of the GUI displays 100 randomly selected rows of the single-cell data table, which is useful to view the table along with the applied modifications.

## Conclusions

High throughput high content image analysis of single-cell data leads to large amounts of quantitative data. H5CellProfiler is a user-friendly solution to analyze such data without the requirement of a database. The current implementation of H5CellProfiler is based on the HDF5 output of the popular HCI image analysis tool CellProfiler. The benefit of such an implementation, in contrast to storing the data in a database, is that the entire pipeline can be performed on a single desktop without the need to set up a database and client software.

The current data workflow of CellProfiler is based on exporting the data to a database followed by analysis with CellProfiler Analyst (Jones et al., 2008) which is based on supervised single-cell machine-learning approaches. Additionally, the CellProfiler data can be exported in the form of spreadsheets, however, this quickly becomes cumbersome in case of large datasets.

We propose the use of the image analysis and data analysis workflow with the use of CellProfiler and R package CellProfiler for the biologist who prefers to handle their own data analysis and graphics. This will save time in the experimental design to the final displayed results cycle and avoids communication difficulties between biologists and computational scientists.

For the more computational adherent scientists — H5CellProfiler has the additional benefit of being based on R which is a highly suited platform for data analysis, statistics, and graphics.

## Code availability

H5CellProfiler is available at https://doi.org/10.5281/zenodo.5484432(Wink and Burger, 2021).

### Basic code description

User-selected data from HDF5 together with the user-provided metadata file is organized into a data.table object with one row per parent object and all associated variables as columns. The output is an `outputList.Rdata` file containing one or several list objects in the case of multiple HDF5 files. The first list object contains the data.table, metadata, user arguments, and a default summary data.table. The `outputList.Rdata` file is used as the input for the `server.R` and `ui.R` code from the Shiny application for the graphical user interface.

Argument class, string length, object definitions, and relations using regular expressions matching to HDF5 paths (of `h5ls` of `$group`, see the rhdf5 vignette for details) are checked and error messages are thrown to inform the user on the type of mistake.

The timestamp of the CellProfiler HDF5 output is extracted with a regular expression. The tracked object and possibly track distance are extracted from the HDF5 paths and in addition, the following track-related paths are defined: tracking label, track parent object numbers, x and y coordinates of tracked object and distance traveled. These track-related HDF5 paths are added to the user-defined paths of `myFeaturesPathsA`.

The `hdf5IndexFun` function pulls data from the HDF5 file and has parameters `hdf5path, dataname`, and `rowIndName`. Parameter `hdf5path` points towards the location of the data in the HDF5 file, `dataname` and `rowIndName` represent the name of the data inside the object and the name of the object index data of that object, respectively. To understand how `hdf5IndexFun` operates, it is important to understand the high-level data organization.

Each object in the HDF5 file has information blocks, for example, the measurements and metadata. It’s object numbers and, if applicable, child or parent object numbers (Fig. 6). When pulled from the HDF5 file by the `h5read` function of the rhdf5 package into memory, each information block of a certain object is represented as a list containing a 1 *×* n objects data array and a 3 *×* n images index array with n being the number of unique entries of that particular object. The data array contains the measurements and object numbers or relation-specific numbers, the index array contains the image number (first column) and the range of rows within the data array to which this image number belongs (2nd and 3rd columns). To find out which object number a particular set of data points belongs to one must look at the data entries within the `Number_Object_Number` information block. Since a cytoplasm object is a derived secondary object, the parent object number of each object must also be determined to link the measurement to the correct parent object. Hence, depending on the type of object different information about the object is required which explains why the user must provide this information. An alternative solution would be to use regular expressions to search for the ‘Parent’ and or ‘Child’ strings to automatically determine these relations. Note that the order of the measurements and annotation information of the objects within an image is constant, so the data of the image object, parent object, and child objects can be gathered per image at a time and then merged using the extracted image numbers, object numbers and parent object numbers. The `hdf5IndFun` function reformats the data using the index information (the index array has an entry for each image) by unstacking the row ranges (which points to the rows in the data array) and these are matched to the row number of the data array. The output is a ‘data.table’ object for a single measurement/annotation from a single object containing the data array entries, the image numbers, and the row index numbers. The row index numbers are not needed after the merge operation of the data and index arrays because the data.tables for each measurement of that same object will be identical. In addition, merging the data.tables of the different objects will depend on the extracted data arrays containing the object numbers of the parent objects and the parent object numbers of the child objects.

**Figure 6:**
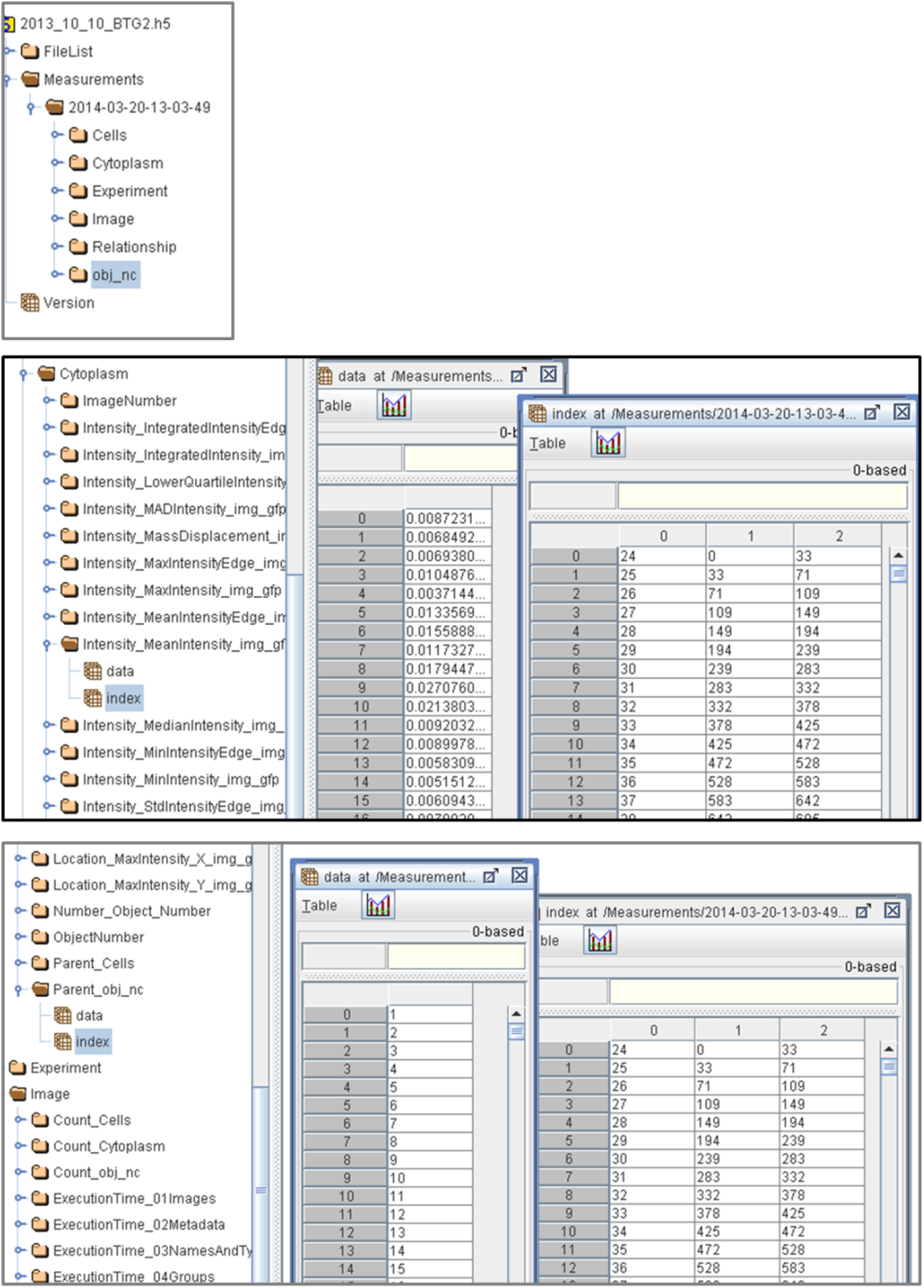
Example of the data organization in an HDF5 file. The top panel shows object blocks Cells, Cytoplasm, Experiment, Image, Relationship, and obj nc. The middle panel displays a level deeper into the object blocks, for example, the information blocks containing measurement and annotation information. The bottom panel shows that object numbers or parent object numbers can be linked when a different information block is included.

